## Supplementary Figures for "Highly sensitive spatial transcriptomics using FISHnCHIPs of multiple co-expressed genes"

### Extended Data Figure 1

**a**

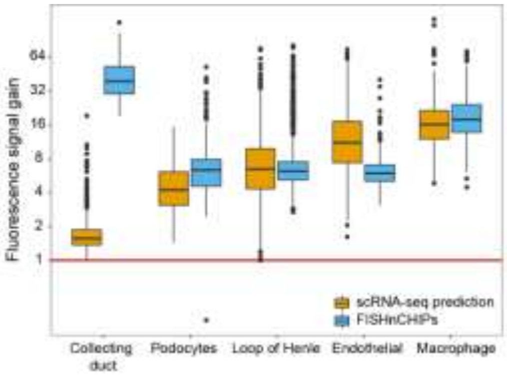


**b**

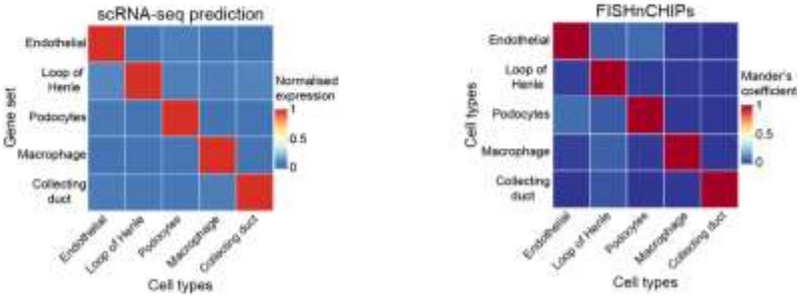


**Extended Data Fig. 1 Quantification of FISHnCHIPs (cell-centric) signal and specificity in the mouse kidney (related to Fig. 2).**

(**a**) Box plots of the ratio of mean fluorescence intensity per cell of FISHnCHIPs to smFISH (blue), and scRNA-seq predictions: the ratio of counts for 14-23 genes to the top DE gene (yellow). Number of cells (FISHnCHIPs): collecting duct: 146, podocytes: 461, loop of Henle: 727, endothelial: 400, macrophage: 341. Number of cells (scRNA-seq): collecting duct: 1,825, podocytes: 77, loop of Henle: 1,496, endothelial: 701, macrophage: 216. The box plots show the median (center line), the first and third quartiles (box limits), and 1.5× the interquartile range (whiskers). Red line indicates where the fluorescence signal gain is 1. (**b**) Predicted signal crosstalk (left heatmap): Normalized mean scRNA-seq counts for FISHnCHIPs genes across the 5 cell types. Number of cells analyzed is the same as in J. Measured FISHnCHIPs crosstalk (right heatmap): Mander's overlap coefficient across the 5 cell-type channels shown in Fig. 2.

### Extended Data Figure 2

**a**

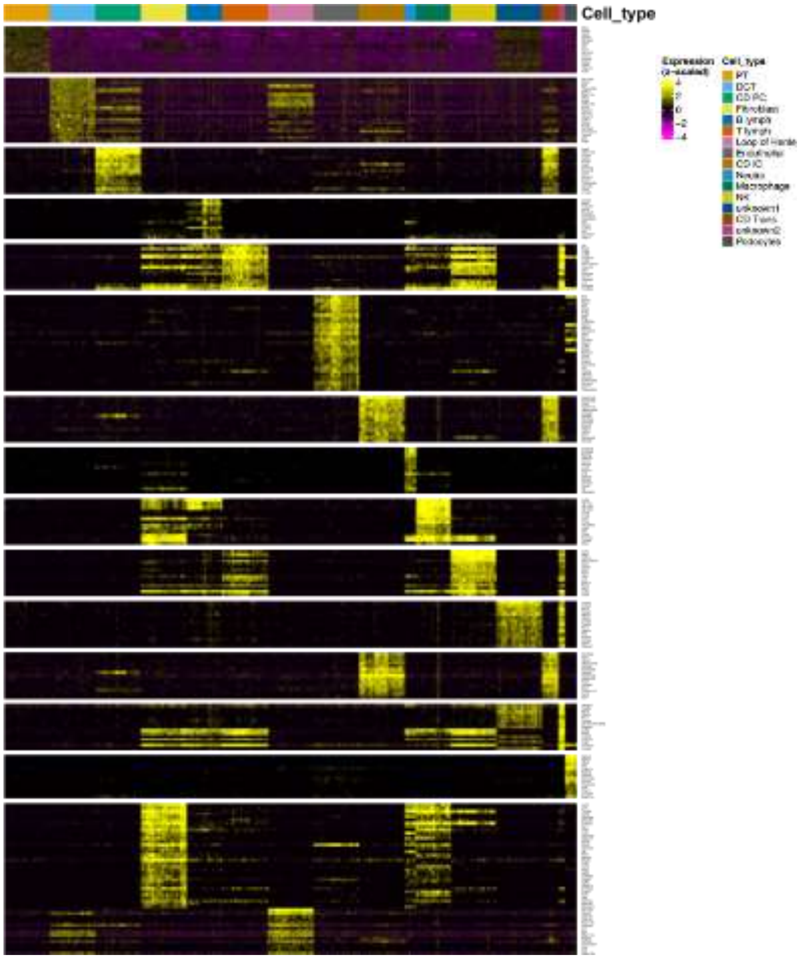


**b**

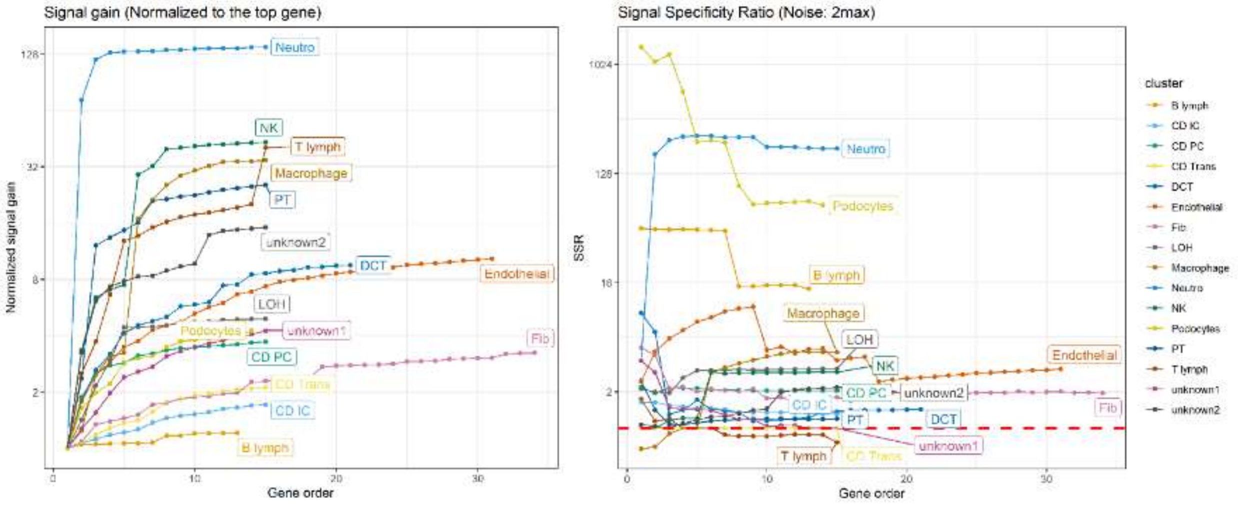

**c**

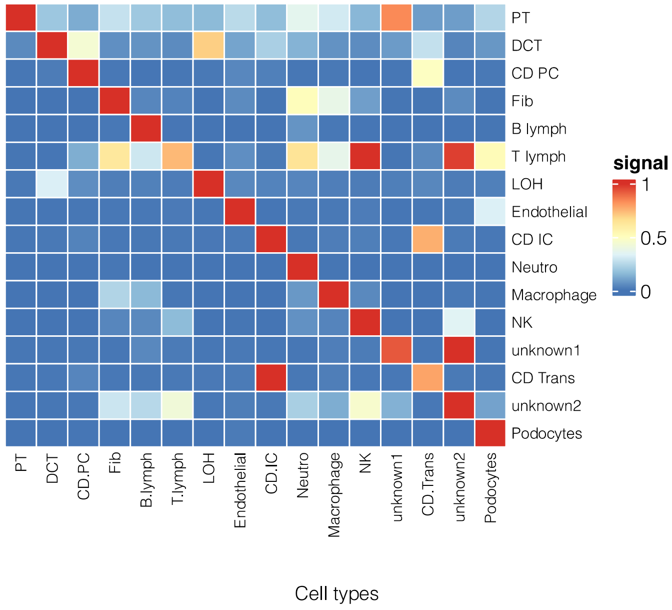


**Extended Data Fig. 2 Computational prediction of FISHnCHIPs (cell-centric) signal gain and specificity.**

(**a**) scRNA-seq[25] gene expression heatmap of a FISHnCHIPs gene panel targeting all the previously annotated mouse kidney cell types, sampling a maximum of 300 cells per cluster. (**b**) Predicted Signal Gain (SG) and Signal Specificity Ratio (SSR) as a function of the number of FISHnCHIPs genes. We defined SG as the ratio of the sum of counts for FISHnCHIPs genes to that of the top DE gene, and SSR as the ratio of the sum of counts for FISHnCHIPs genes in the target cell type to that in the most likely off-target cell type. When SSR approaches unity, the fluorescence intensity for the cell type of interest will be equal to an off-target cell type, rendering them indistinguishable. (**c**) Predicted signal crosstalk: Heatmap of the normalized mean scRNA-seq counts of the FISHnCHIPs gene panel across all kidney cell types.

### Extended Data Figure 3

**a**

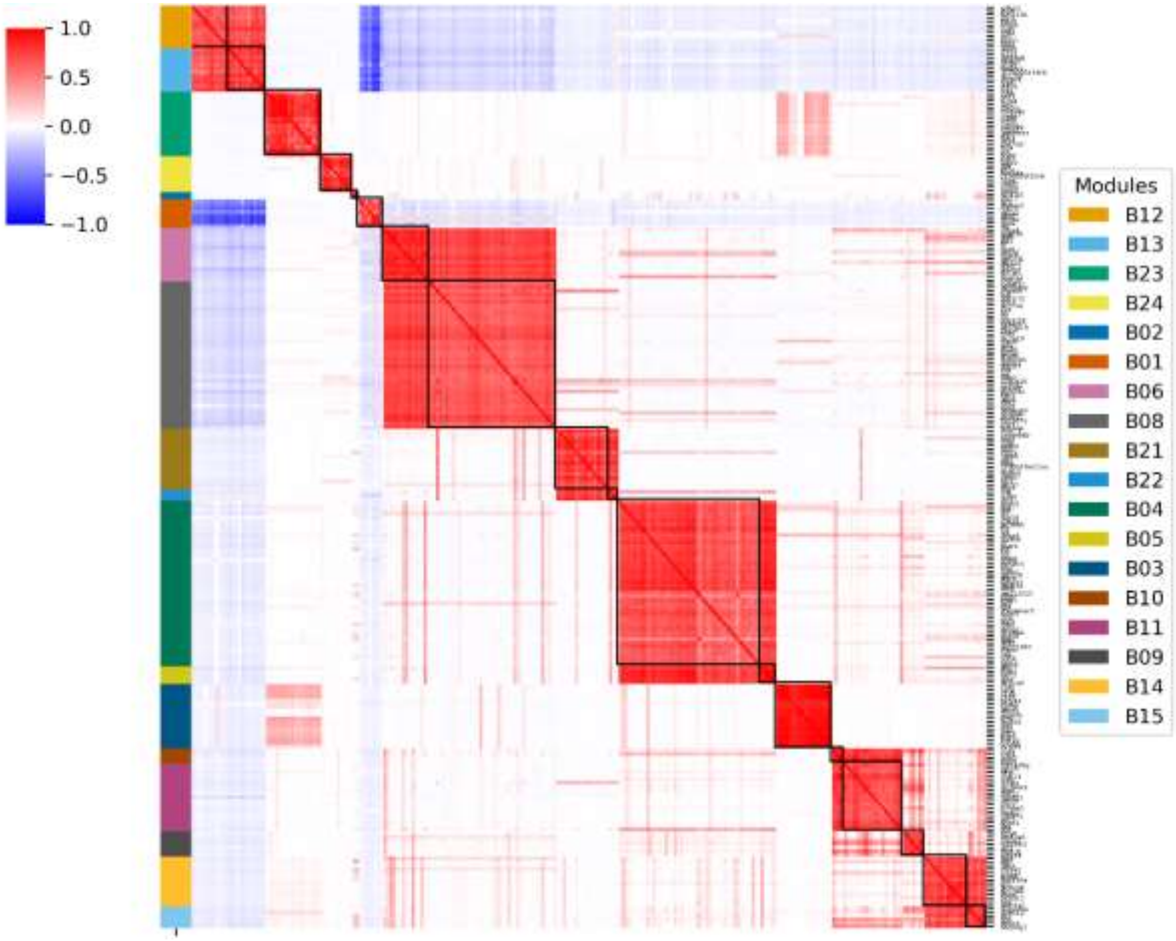

**b**

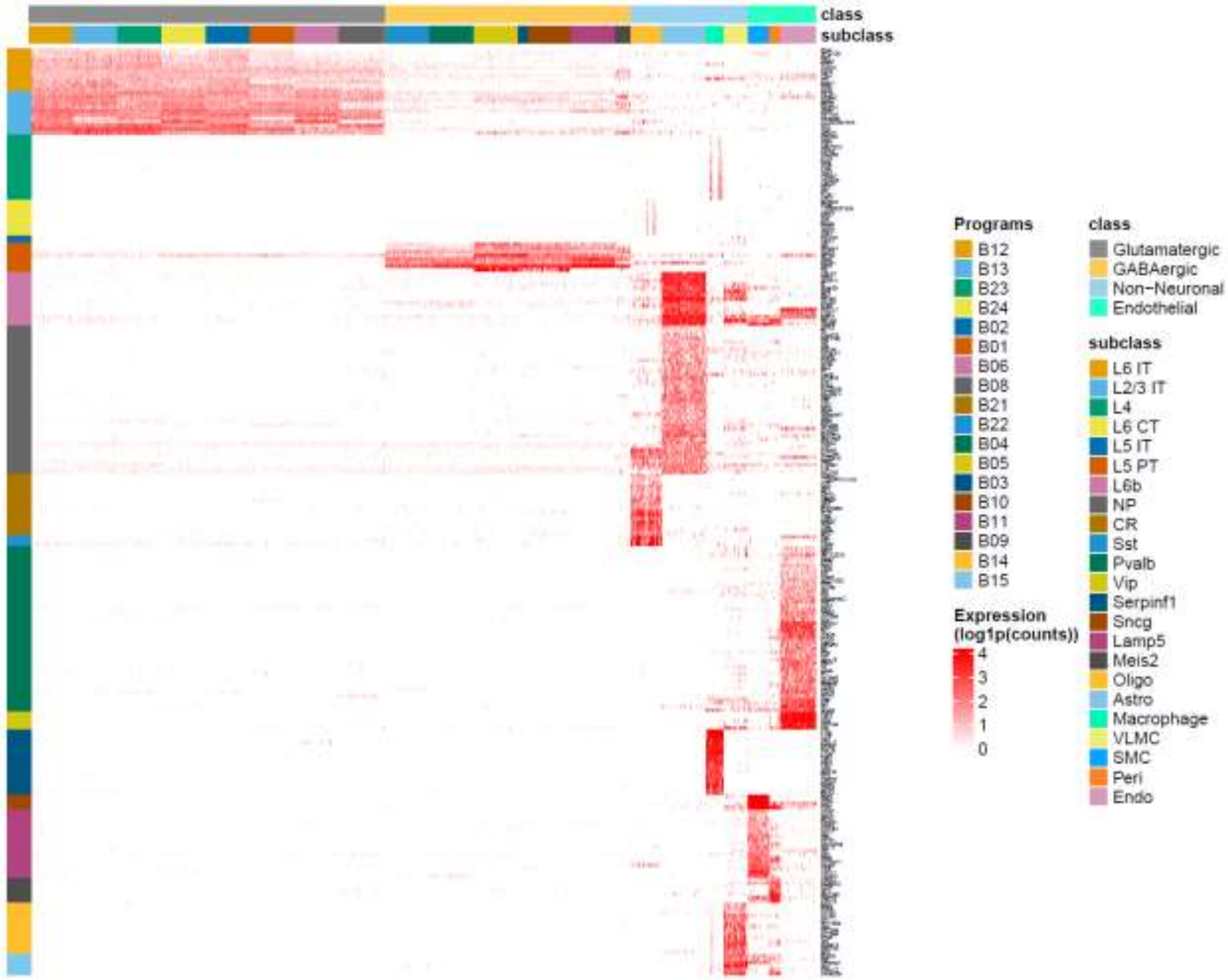
class
subclass
Programs
B12
B13
B23
B24
B02
B01
B06
B08
B21
B22
B04
B05
B03
B10
B11
B09
B14
B15
Expression
(log1p(counts))
4
3
2
1
0
class
Glutamatergic
GABAergic
Non−Neuronal
Endothelial
subclass
L6 IT
L2/3 IT
L4
L6 CT
L5 IT
L5 PT
L6b
NP
CR
Sst
Pvalb
Vip
Serpinf1
Sncg
Lamp5
Meis2
Oligo
Astro
Macrophage
VLMC
SMC
Peri
Endo

**c**

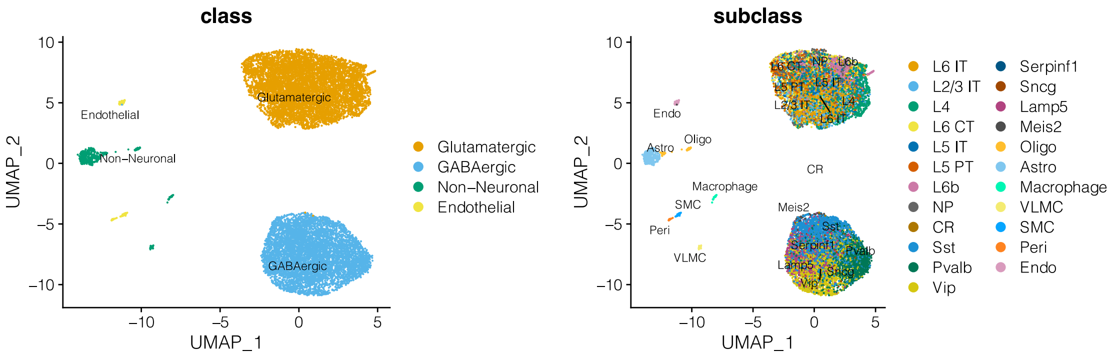
class
Endothelial
Non−Neuronal
Glutamatergic
GABAergic
UMAP_2
UMAP_1
Glutamatergic
GABAergic
Non−Neuronal
Endothelial
subclass
Endo
Astro
Oligo
CR
Macrophage
SMC
Peri
VLMC
Meis2
UMAP_2
UMAP_1
L6 IT
L2/3 IT
L4
L6 CT
L5 IT
L5 PT
L6b
NP
CR
Sst
Pvalb
Vip
Serpinf1
Sncg
Lamp5
Meis2
Oligo
Astro
Macrophage
VLMC
SMC
Peri
Endo

**d**

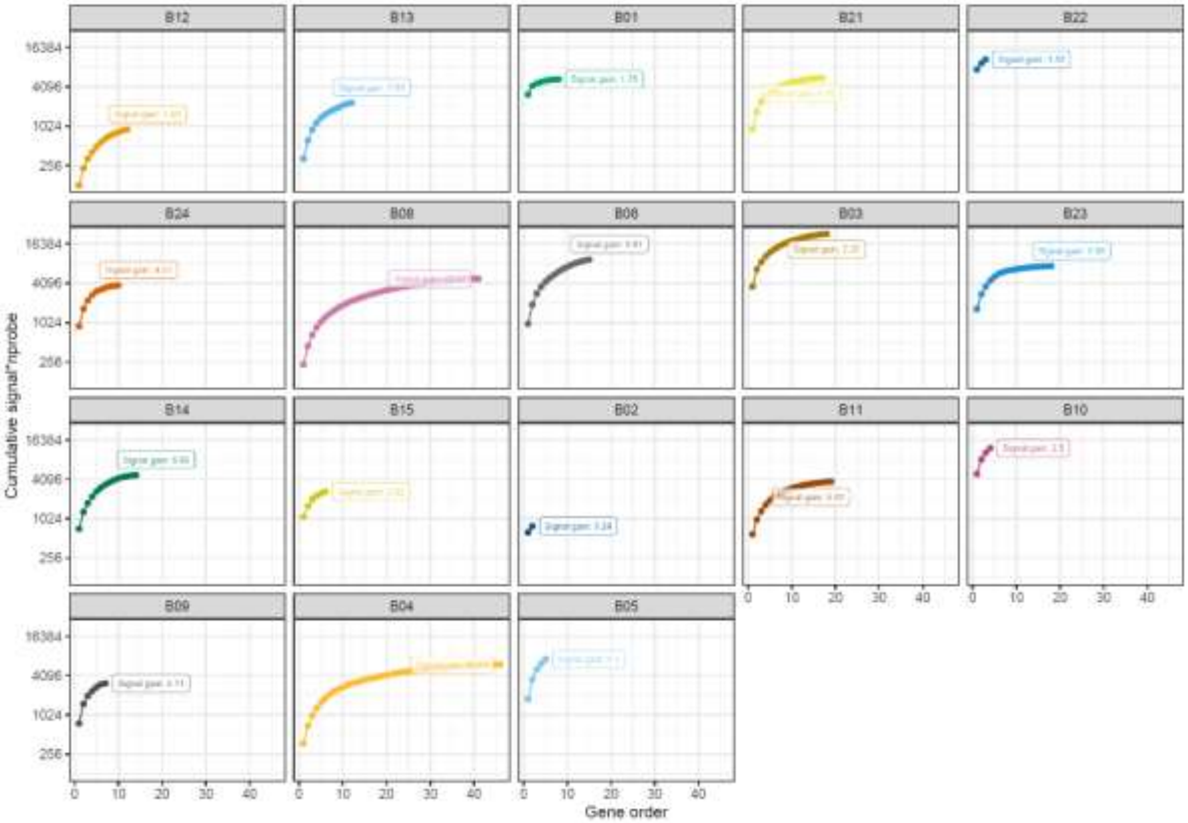

B12
B13
B01
B21
B22
B24
B08
B06
B03
B23
B14
B15
B02
B11
B10
B09
B04
B05
16384
4096
1024
256
Cumulative signal*nprobe
0
10
20
30
40
Gene order


**e**

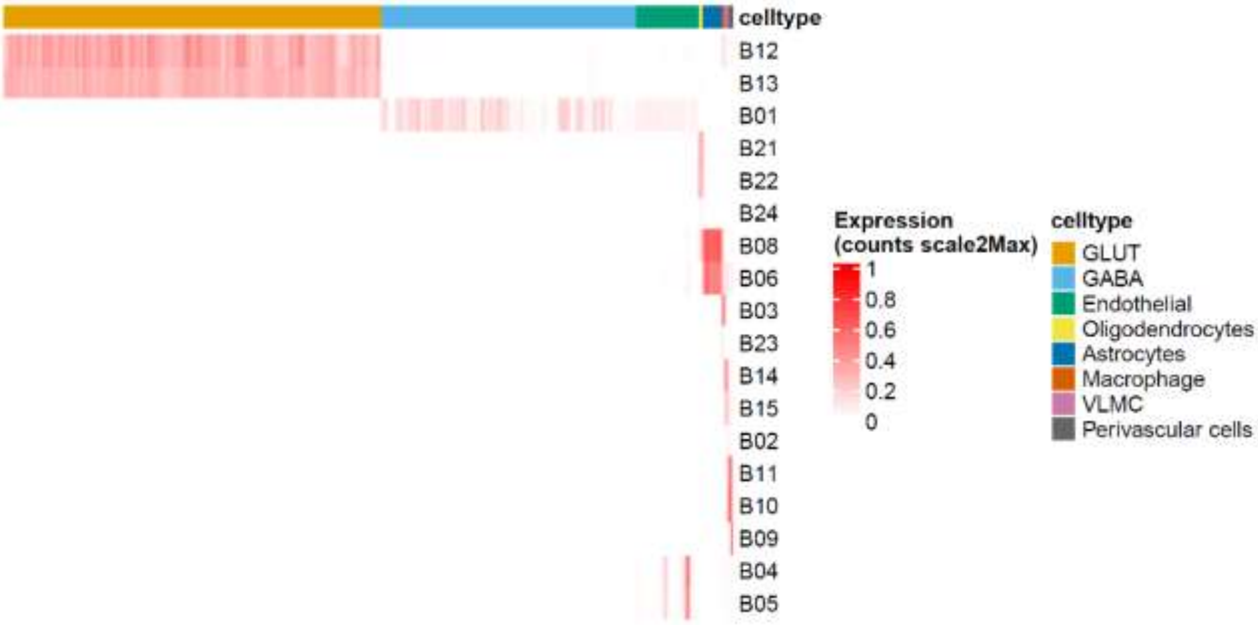

celltype
B12
B13
B01
B21
B22
B24
B08
B06
B03
B23
B14
B15
B02
B11
B10
B09
B04
B05
Expression
(counts scale2Max)
1
0.8
0.6
0.4
0.2
0
celltype
GLUT
GABA
Endothelial
Oligodendrocytes
Astrocytes
Macrophage
VLMC
Perivascular cells

**Extended Data Fig. 3 Gene module based FISHnCHIPs library design for the mouse cortex library (imaged in Fig. 3).**

(**a**) scRNA-seq[27] gene-gene correlation heatmap (with gene names) for the 255 feature genes in the mouse cortex library (imaged in Fig. 3). We computed their pair-wise Pearson's correlation coefficient and clustered the correlation matrix using the Leiden algorithm. The gene partitions were further sub-clustered using hierarchical clustering into 18 modules. (**b**) scRNA-seq gene expression heatmap for the 255 genes. (**c**) UMAP representation of the clustering of module-cell (meta-gene) expression, indicated by the labels provided by the scRNA-seq reference dataset. ~8 cell types are clearly separated with the selected features. (**d**) Predicted conservative Signal Gain (cumulative), which is defined as the ratio of the panel signal to the highest gene signal, as a function of the number of genes. (**e**) Predicted module-cell expression heatmap. We grouped the cluster labels into the 8 resolvable cell types (see Supplementary Table 5).

### Extended Data Figure 4

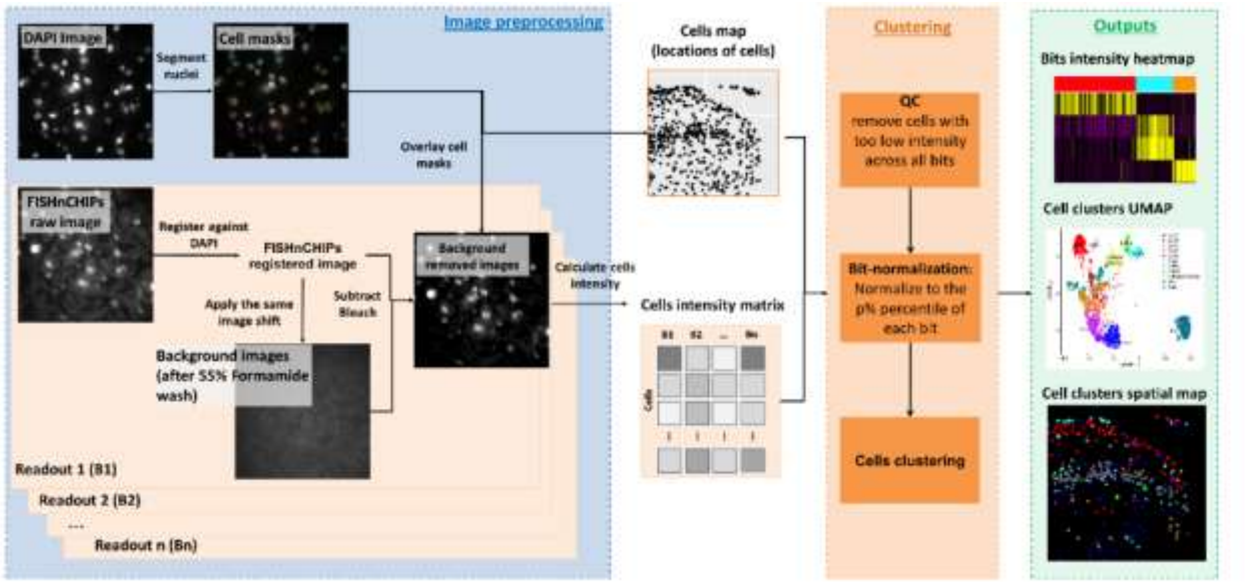


**Extended Data Fig. 4 Overview of the FISHnCHIPs image processing and data analysis workflow.**

Stepwise data processing (see methods, as well as supplementary software): Inputs are DAPI, FISHnCHIPs, and background (after 55% formamide wash) images. 1) Preprocessing steps include segmentation based on DAPI images to generate cell masks; 2) Registration and background subtraction of FISHnCHIPs images; 3) Cell masks are used to generate cell intensity matrix with a list of cell centroids. 4) Clustering of the cell intensity matrix. 5) Outputs can be visualized in a heatmap, UMAP, or spatial map. Outputs can also be subjected to further analyses, such as classifications of spatial patterns and analysis of cell-cell interactions.

### Extended Data Figure 5

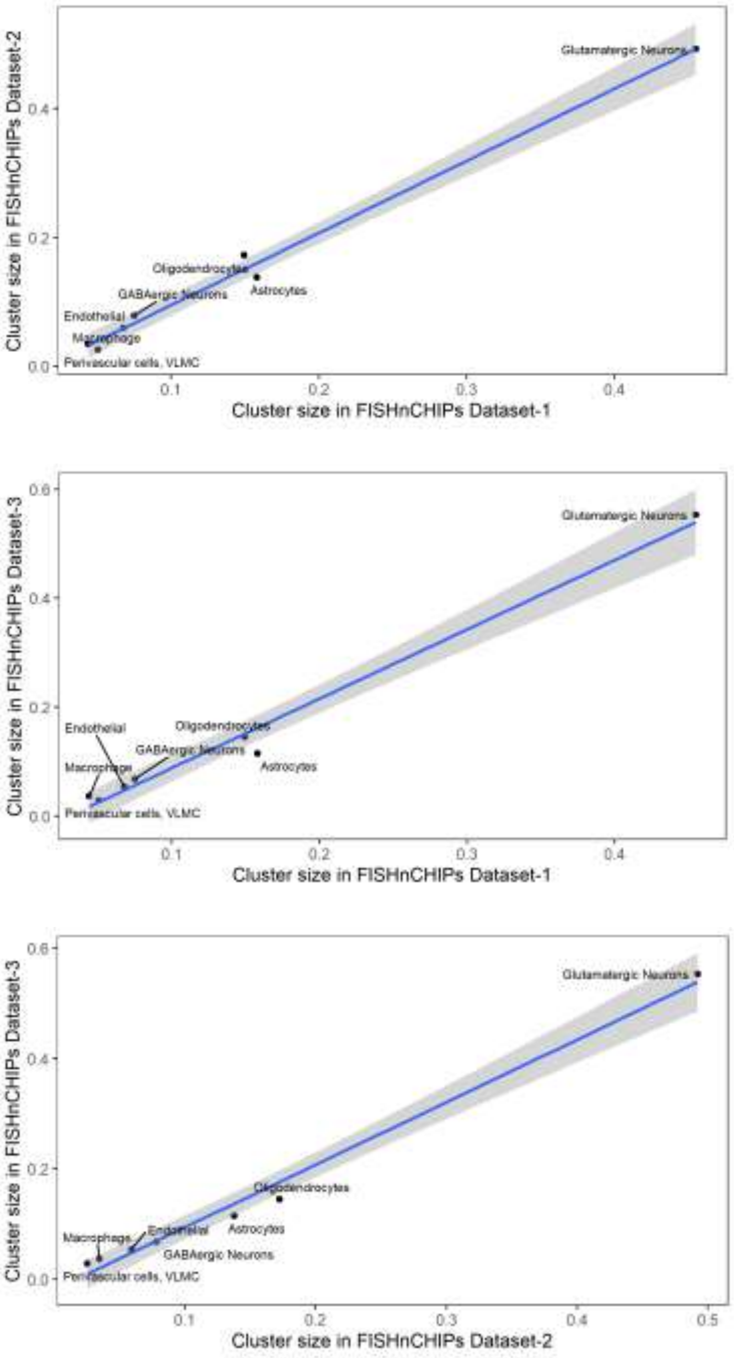


**Extended Data Fig. 5 Reproducibility of the mouse brain FISHnCHIPs data (related to Fig. 3) among technical triplicates.**

Scatter plots of cell type abundances between technical replicates.

### Extended Data Figure 6

**a**

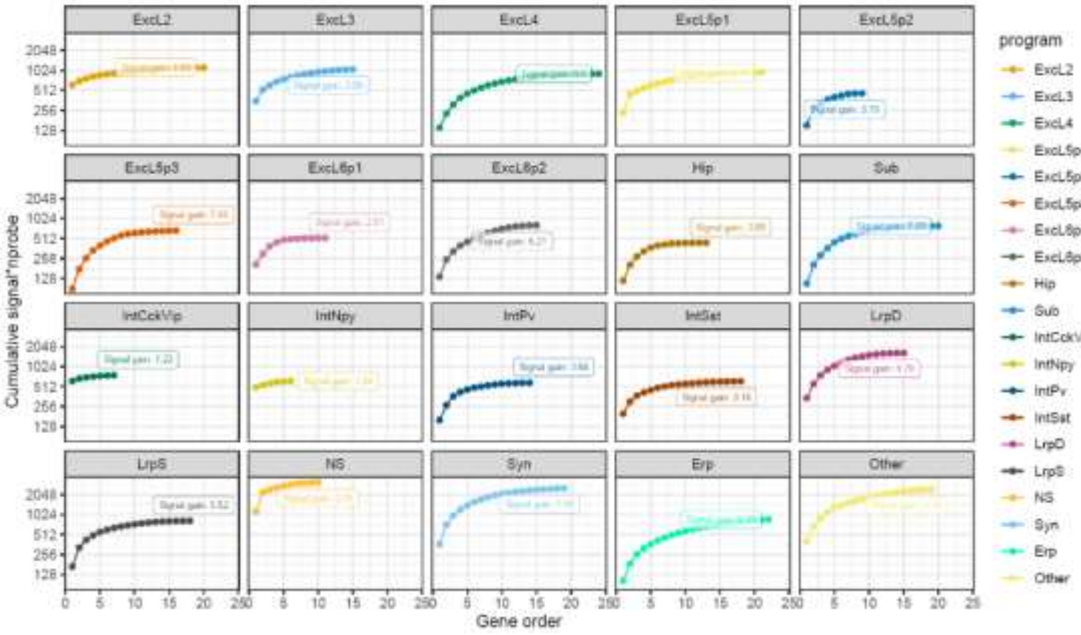


**b**

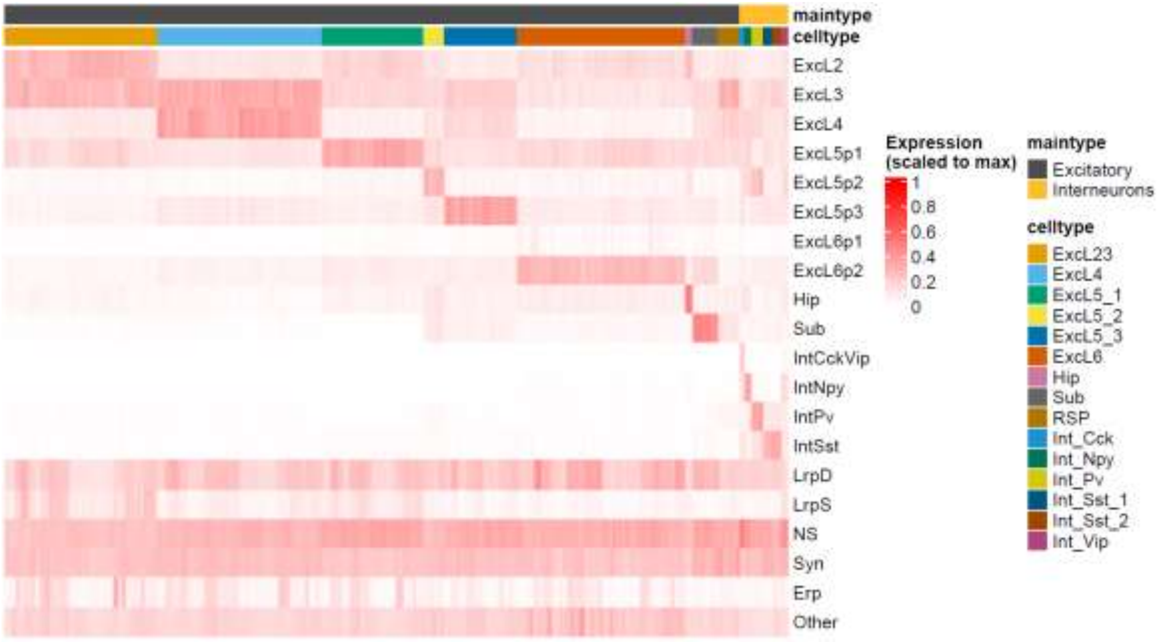

**c**

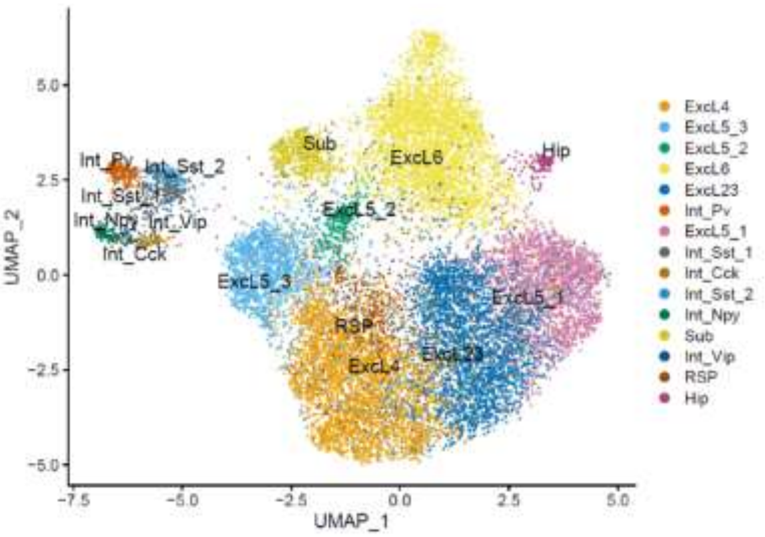
UMAP_2
UMAP_1
Int_Pv
Int_Sst_2
Int_Sst_1
Int_Npy
Int_Vip
Int_Cck
Sub
ExcL6
Hip
ExcL5_2
ExcL5_3
RSP
ExcL4
ExcL23
ExcL5_1
ExcL4
ExcL5_3
ExcL5_2
ExcL6
ExcL23
Int_Pv
ExcL5_1
Int_Sst_1
Int_Cck
Int_Sst_2
Int_Npy
Sub
Int_Vip
RSP
Hip

**d**

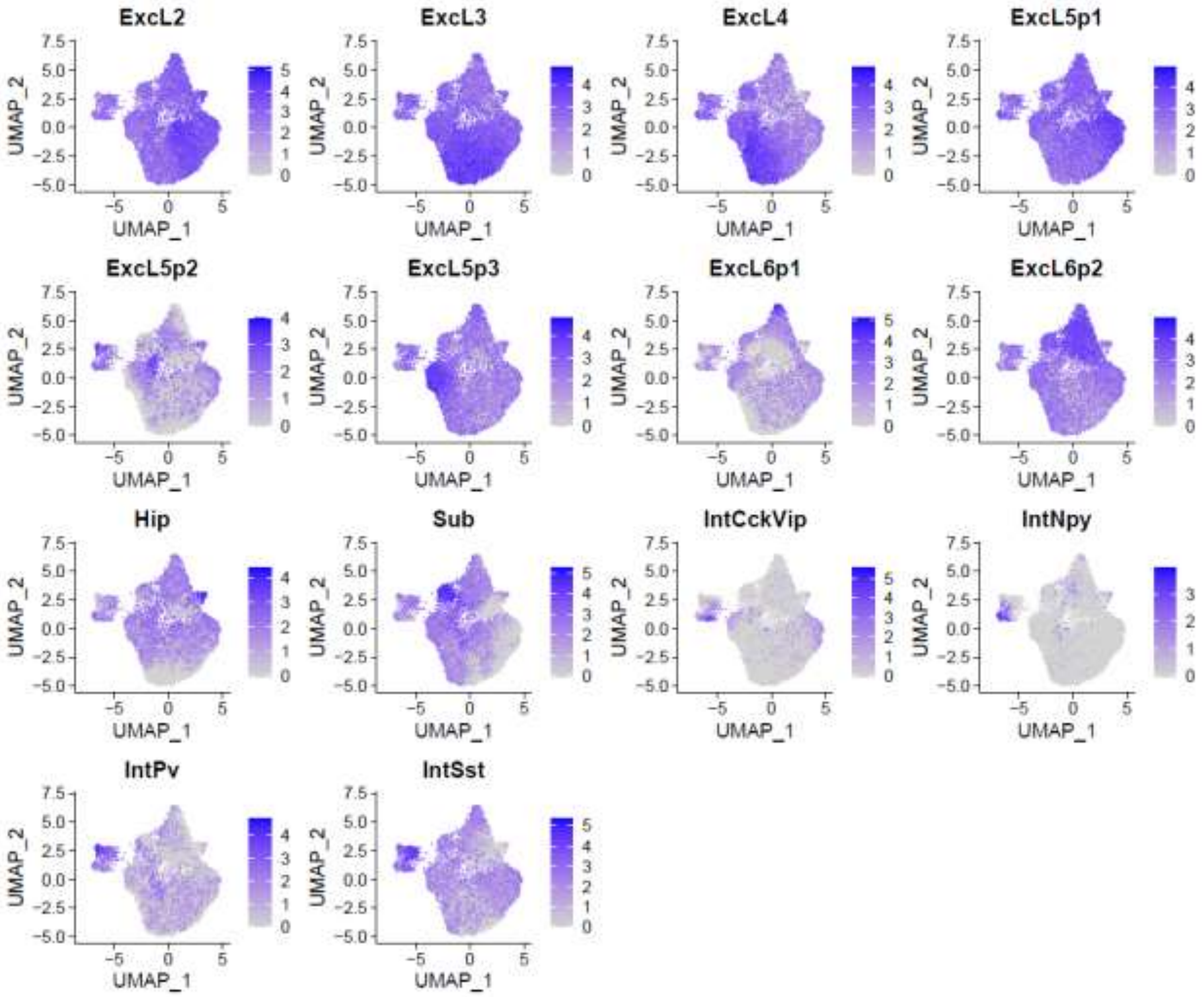
ExcL2
ExcL3
ExcL4
ExcL5p1
ExcL5p2
ExcL5p3
ExcL6p1
ExcL6p2
Hip
Sub
IntCckVip
IntNpy
IntPv
IntSst
UMAP_1
UMAP_2

**Extended Data Fig. 6 Evaluation of the gene expression programs based FISHnCHIPs library for the mouse visual cortex (imaged in Fig. 4).**

(**a**) Predicted conservative Signal Gain (cumulative), which is defined as the ratio of the panel signal to the highest gene signal, as a function of the number of genes. (**b**) Predicted Signal Specificity: scRNA-seq expression heatmap for the 20 programs. (**c**) UMAP representation of the clustering of program-cell (meta-gene) expression, indicated by the labels provided by the scRNA-seq reference dataset[59]. (**d**) scRNA-seq feature plots of the 14 identify programs.

### Extended Data Figure 7

**a**

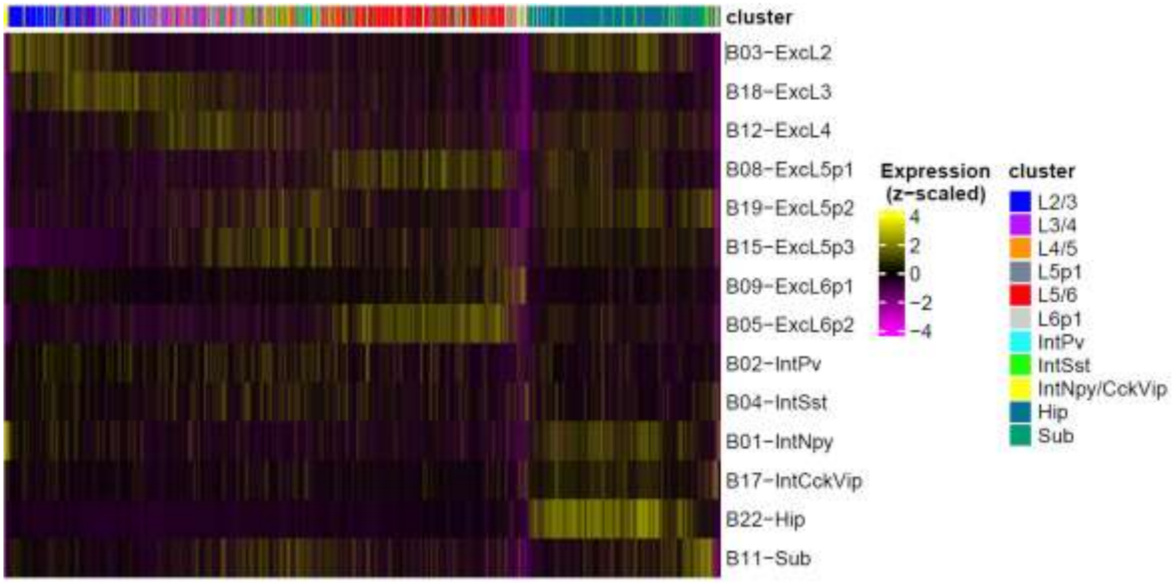


**b**

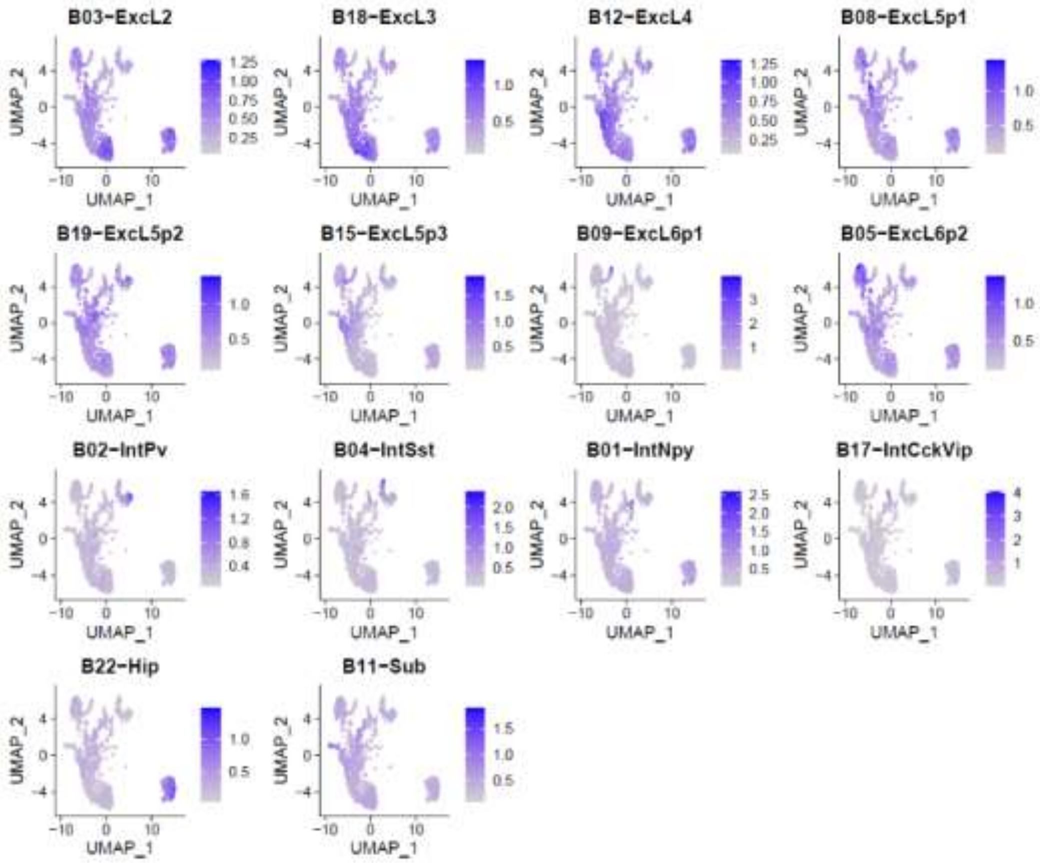

**Extended Data Fig. 7 Gradients of gene expression along the cortical depth of the mouse visual cortex as imaged by FISHnCHIPs (related to Fig. 4).**

(**a**) FISHnCHIPs expression heatmap of the cell-by-program-intensity matrix (cells are ordered by their distance to the outer edge). (**b**) FISHnCHIPs feature plots of the 14 identity programs.

### Extended Data Figure 8

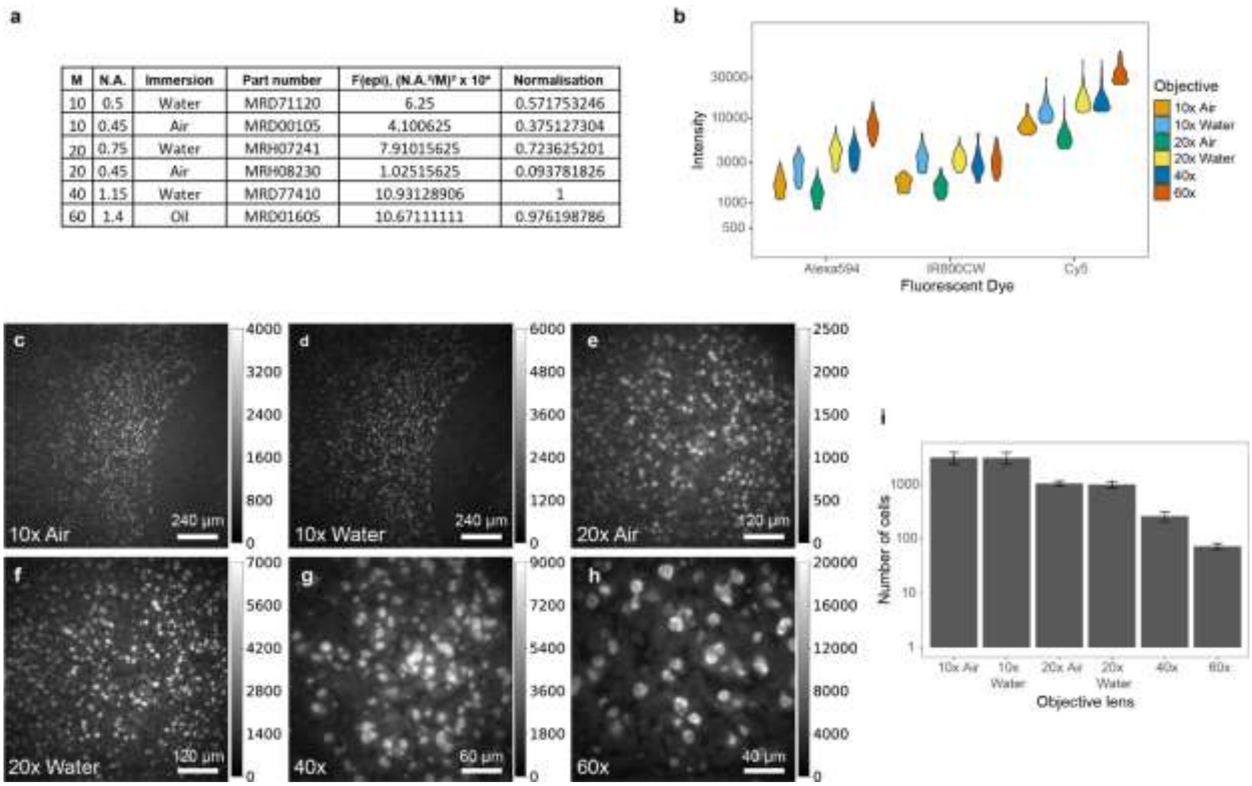

| M | N.A. | Immersion | Part number | $F(epi), (N.A.^4/M)^2 \times 10^4$ | Normalisation |
|---|---|---|---|---|---|
| 10 | 0.5 | Water | MRD71120 | 6.25 | 0.571753246 |
| 10 | 0.45 | Air | MRD00105 | 4.100625 | 0.375127304 |
| 20 | 0.75 | Water | MRH07241 | 7.91015625 | 0.723625201 |
| 20 | 0.45 | Air | MRH08230 | 1.02515625 | 0.093781826 |
| 40 | 1.15 | Water | MRD77410 | 10.93128906 | 1 |
| 60 | 1.4 | Oil | MRD01605 | 10.67111111 | 0.976198786 |



**Extended Data Fig. 8 FISHnCHIPs of the mouse brain imaged under lower magnification.** (**a**) Table of magnification (M), numerical aperture (N.A.), and predicted light gathering power under epi-illumination configuration, F(epi)[60], for 6 different objective lenses. (**b**) Measured mean fluorescence intensity per cell for Alexa594, Cy5, and IR800CW, for 6 different objective lens. (**c-h**) Example unprocessed FISHnCHIPs images (one Field of View, FOV) of the mouse cortex under 6 different objective lenses. Cells labelled with FISHnCHIPs were detected above the background level across all three-color channels, even at the 10x magnification. (**i**) The number of cells detected per FOV (n = 5 FOVs, error bars indicate the standard deviation). Because of the wider field of view, the number of cells imaged was >~40 fold greater when using the 10x versus 60x objective lenses. Average number of cells: 10x air: 3130, 10x water: 3088, 20x air: 1003, 20x water: 1041, 40x: 261, 60x: 73. We chose the 10x water objectives for Fig. 5 data acquisition.

### Extended Data Figure 9

**a**

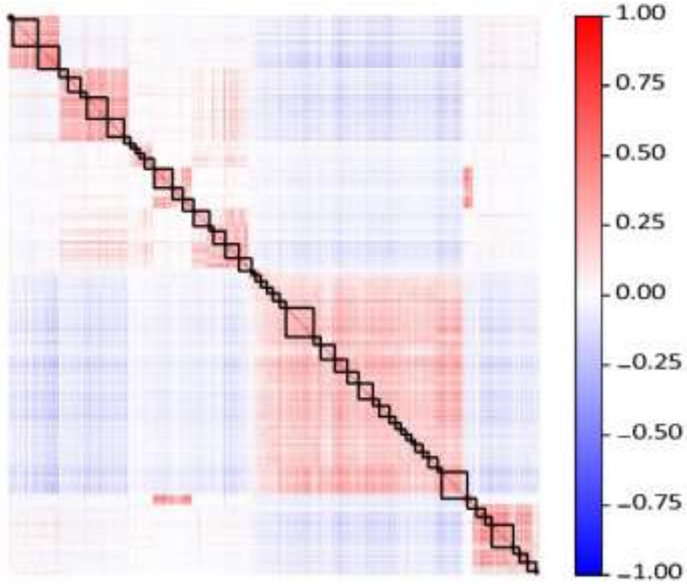


**b**

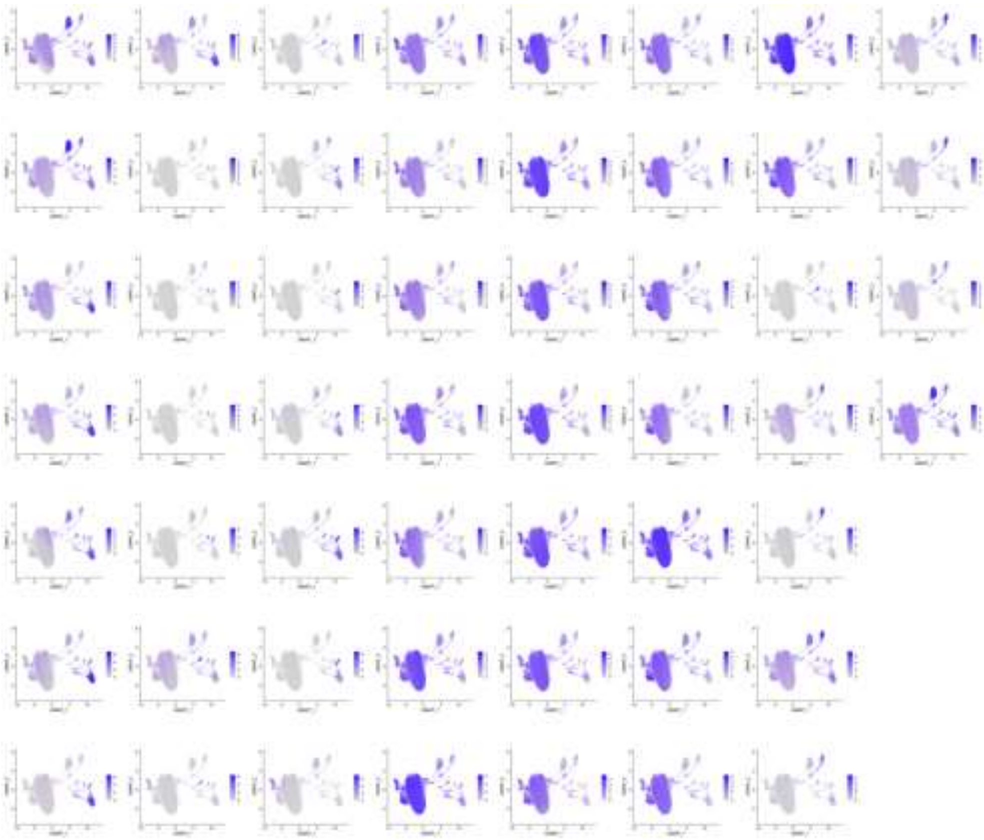

**Extended Data Fig. 9 scRNA-seq gene-gene correlation of the FISHnCHIPs genes targeting 53 gene modules in the mouse cortex (imaged in Fig. 5).**

(**a**) scRNA-seq gene-gene correlation heatmap for the 674 feature genes in the mouse cortex library (imaged in Fig. 5). We computed their pair-wise Pearson's correlation coefficient and clustered the correlation matrix using the Leiden algorithm. The gene partitions were further sub-clustered using hierarchical clustering into 53 gene modules. (**b**) scRNA-seq feature plots of the 53 gene modules.

### Extended Data Figure 10

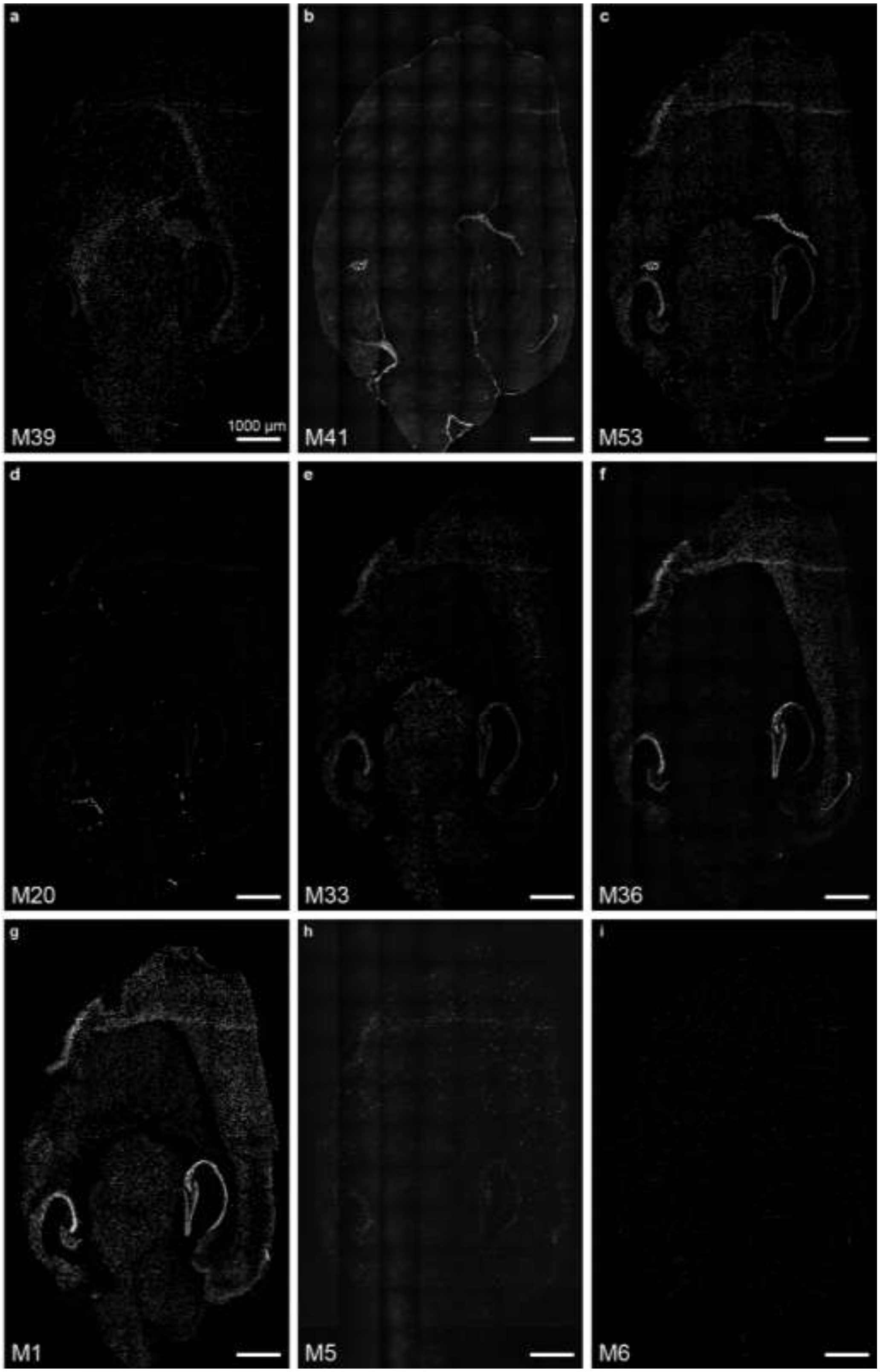

**Extended Data Fig. 10 Example normalized images from the 53-modules FISHnCHIPs profiling** of gene module 39, gene module 41, gene module 53 using Alexa 594 (**a** - **c**), gene module 20, gene module 33, gene module 36 using Cy5 (**d** - **f**), and gene module 1, gene module 5, and gene module 6 using IRDye 800CW (**g** - **i**) under the 10x objective lens. Scale bar, 1000 μm

### Extended Data Figure 11

**a**

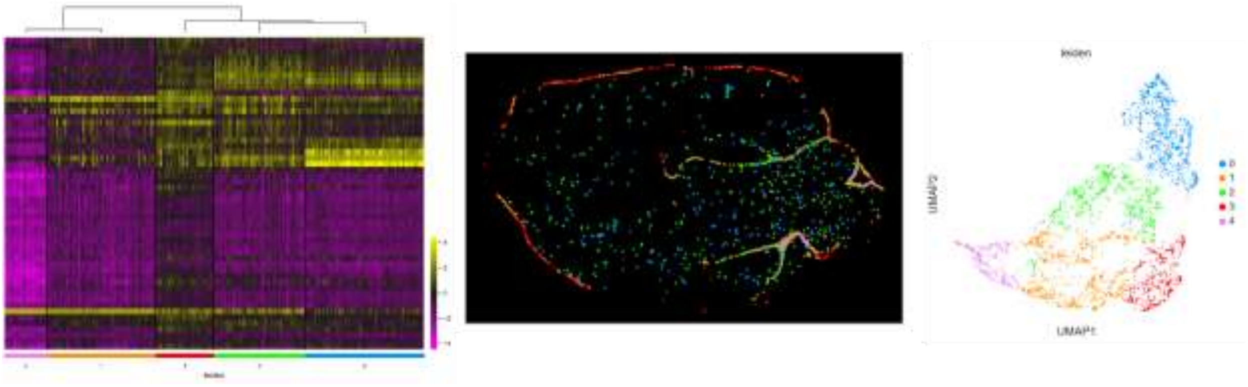


**b**

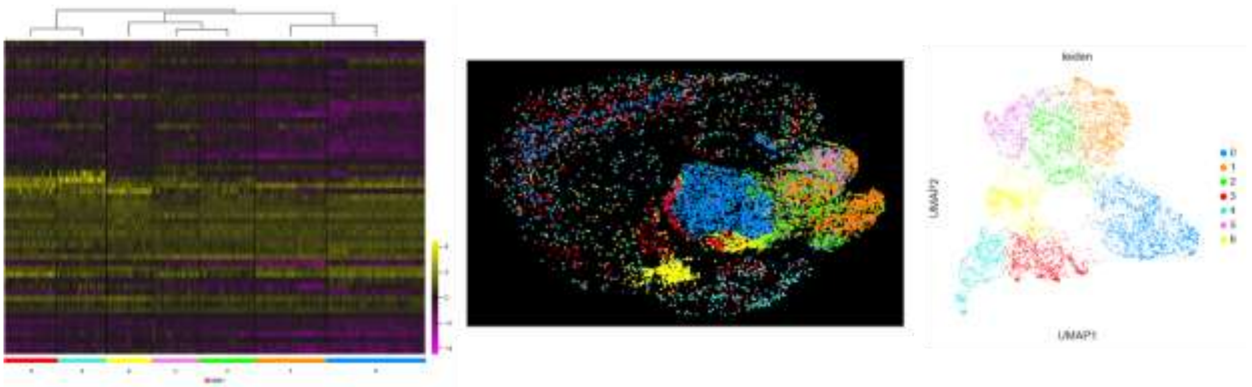


**Extended Data Fig. 11 Sub-clustering analyses of the 53-module FISHnCHIPs data revealed subtypes of blood vessel associated and inhibitory cell subtypes with distinct spatial patterns.** (**a**) FISHnCHIPs expression heatmap, spatial map, and UMAP of the subtypes of blood vessel associated cells (**b**) FISHnCHIPs expression heatmap, spatial map, and UMAP of the subtypes of inhibitory neurons

### Extended Data Figure 12

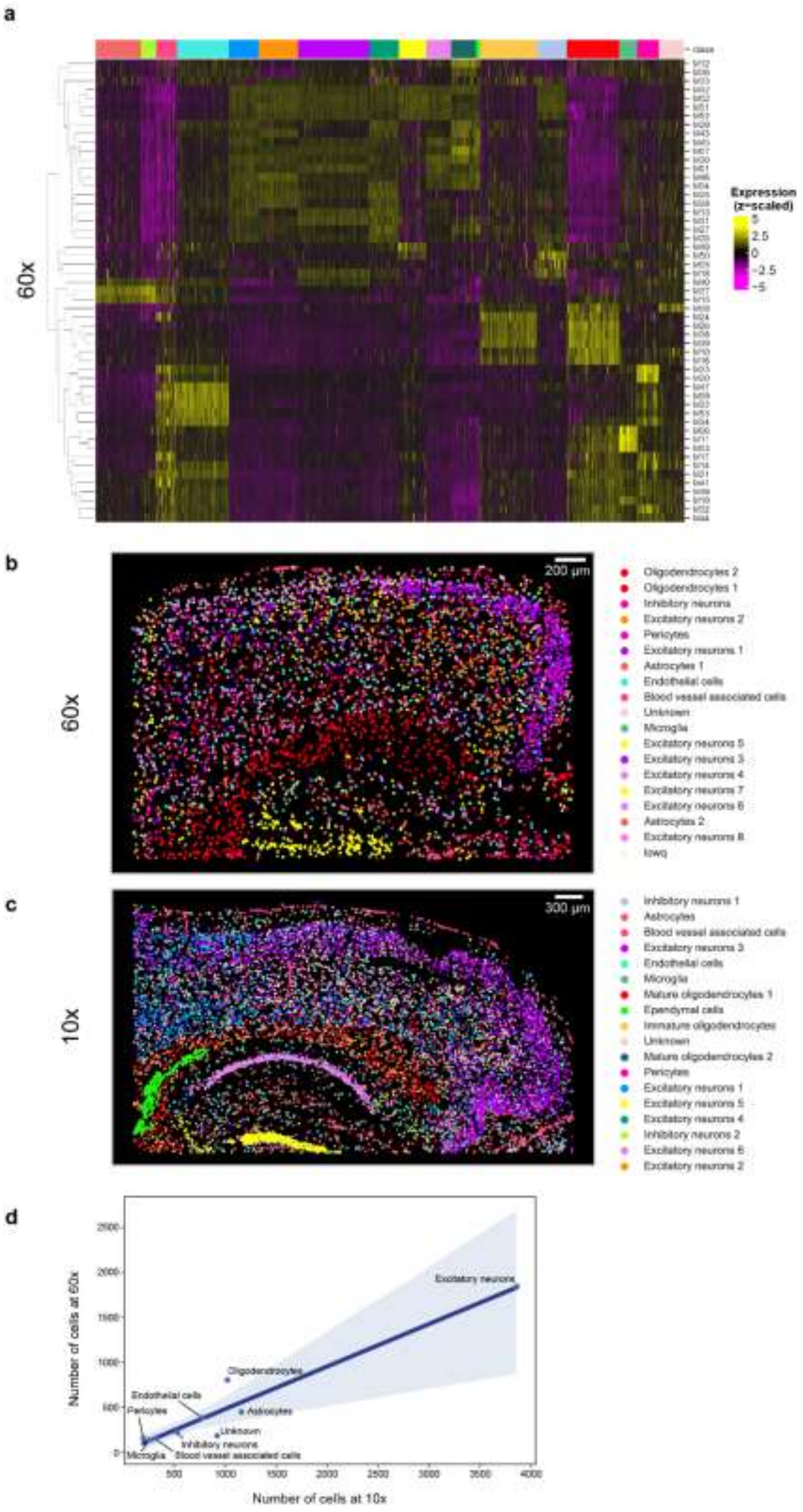

**Extended Data Fig. 12 Proportion of cell types in the 10x and 60x datasets.** (**a**) 53-module FISHnCHIPs expression heatmap for 4,329 cells acquired using the 60x objectives (**b**) Spatial map of FISHnCHIPs cell types (60x) (**c**) 53-module FISHnCHIPs expression heatmap for 9,323 cells acquired using the 10x objectives (Fig. 5 data but cropped to the visual cortex region) (**d**) Scatter plot of cell type frequency detected by 60x versus 10x (cropped to the visual cortex region).

### Extended Data Figure 13

**a**

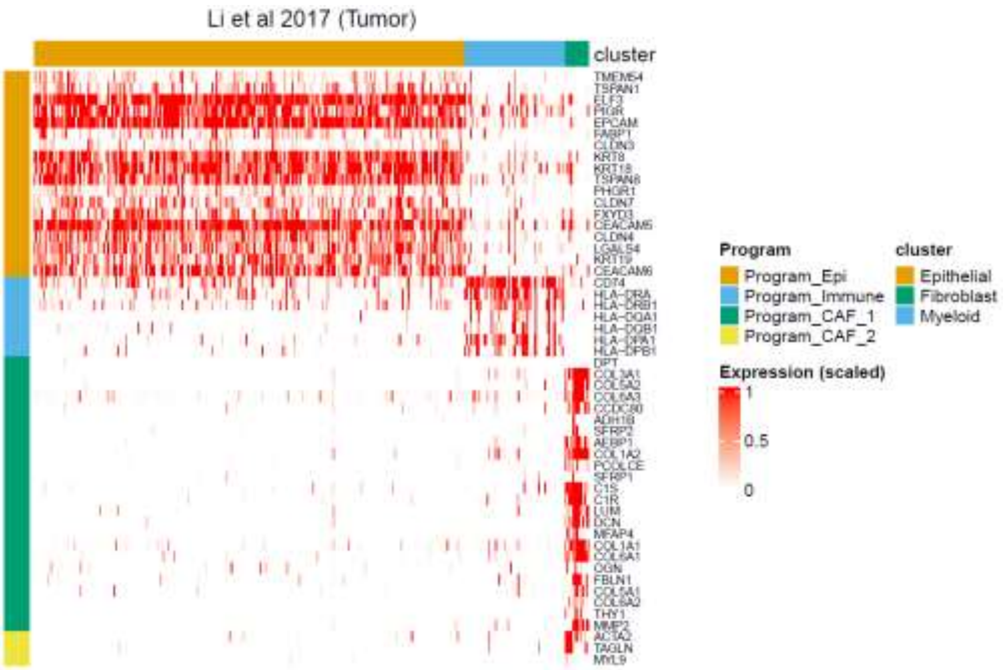


**b**

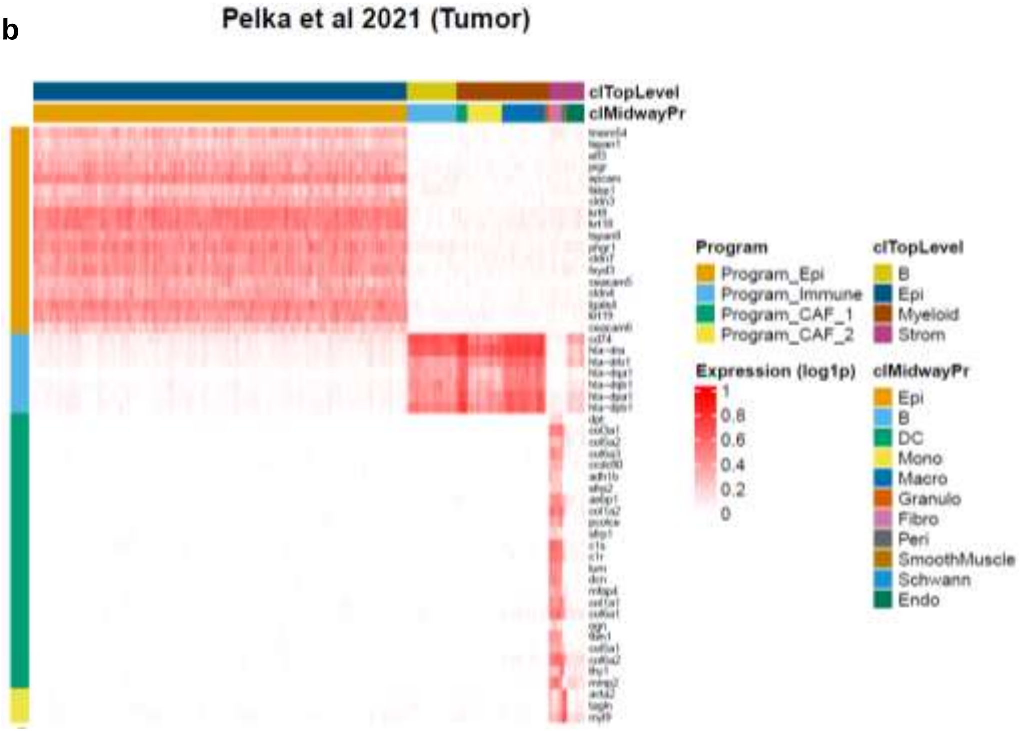


**c**

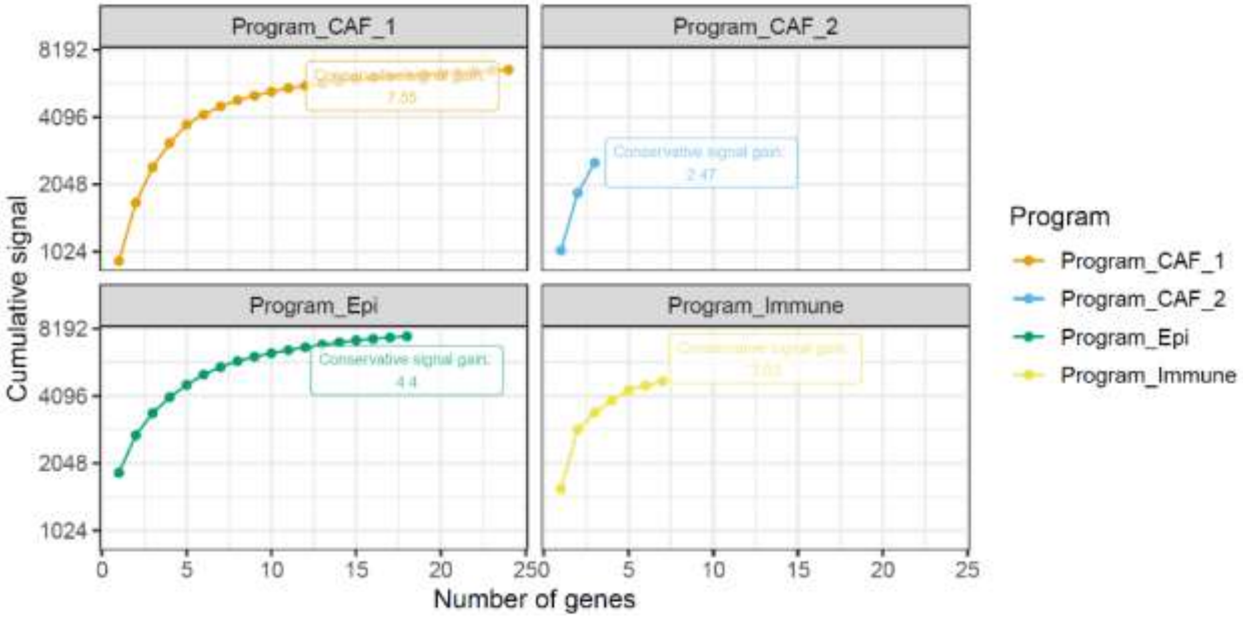


**Extended Data Fig. 13 FISHnCHIPs panel for imaging cancer associated fibroblasts (CAFs) subtypes in human colorectal cancer (CRC) tissue (related to Fig. 6).**

(**a**) scRNA-seq gene expression heatmap of the human CRC FISHnCHIPs panel[34]. (**b**) scRNA-seq gene expression heatmap of the human CRC FISHnCHIPs panel[48]. (**c**) Predicted conservative signal gain for the human CRC FISHnCHIPs panel.

### Extended Data Figure 14

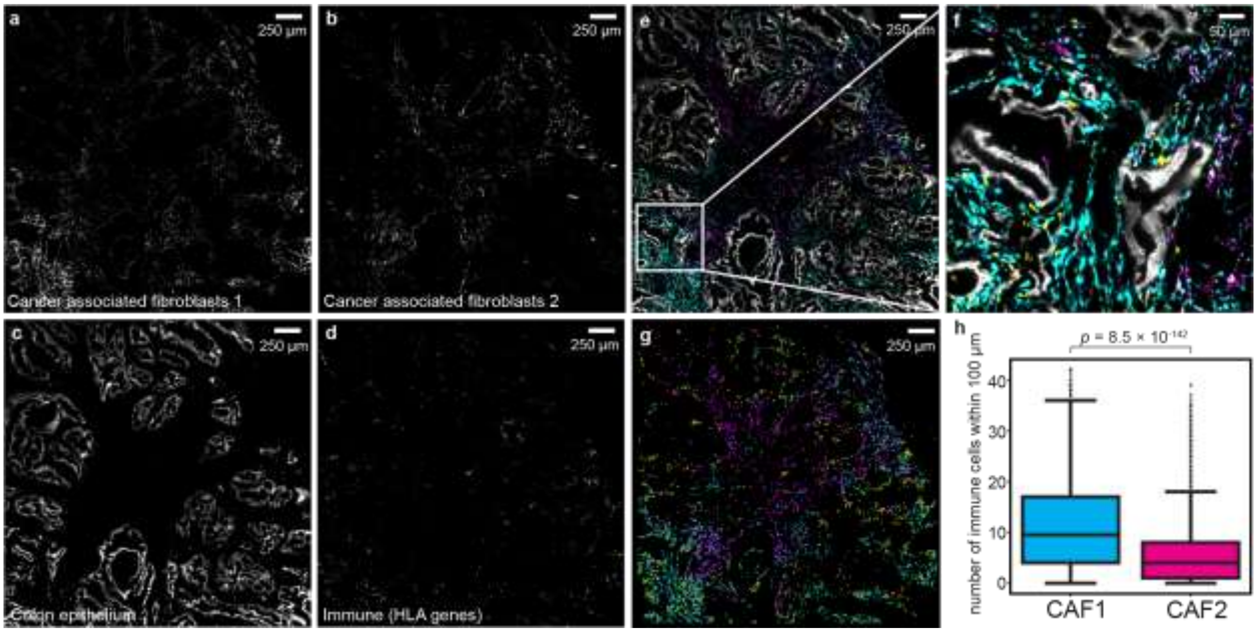


**Extended Data Fig. 14 Technical replicate of FISHnCHIPs on human CRC tissue.**

(**a**) FISHnCHIPs image of CAF-1 subtype. Scale bar, 250 µm.(**b**) FISHnCHIPs image of CAF-2 subtype. Scale bar, 250 µm. (**c**) FISHnCHIPs image of colon epithelium. Scale bar, 250 µm. (**d**) FISHnCHIPs image of immune (HLA genes). Scale bar, 250 µm. (**e**) Composite FISHnCHIPs image. Scale bar, 250 µm. (**f**) Zoom-in of the white box in E. Scale bar, 50 µm. (**g**) Box plots of the number of immune cells within 100 µm radius of CAF-1 (cyan) and CAF-2 (purple) cells. Immune cells were found 0.51-fold less frequently in the vicinity of CAF-2 than CAF-1. Number of cells: CAF-1: 2,548, CAF-2: 2,199. The box plots show the median (centre line), the first and third quartiles (box limits), and 1.5× the interquartile range (whiskers). $p = 8.5 \times 10^{-142}$, 2-sided Mann-Whitney U test.

### Extended Data Figure 15

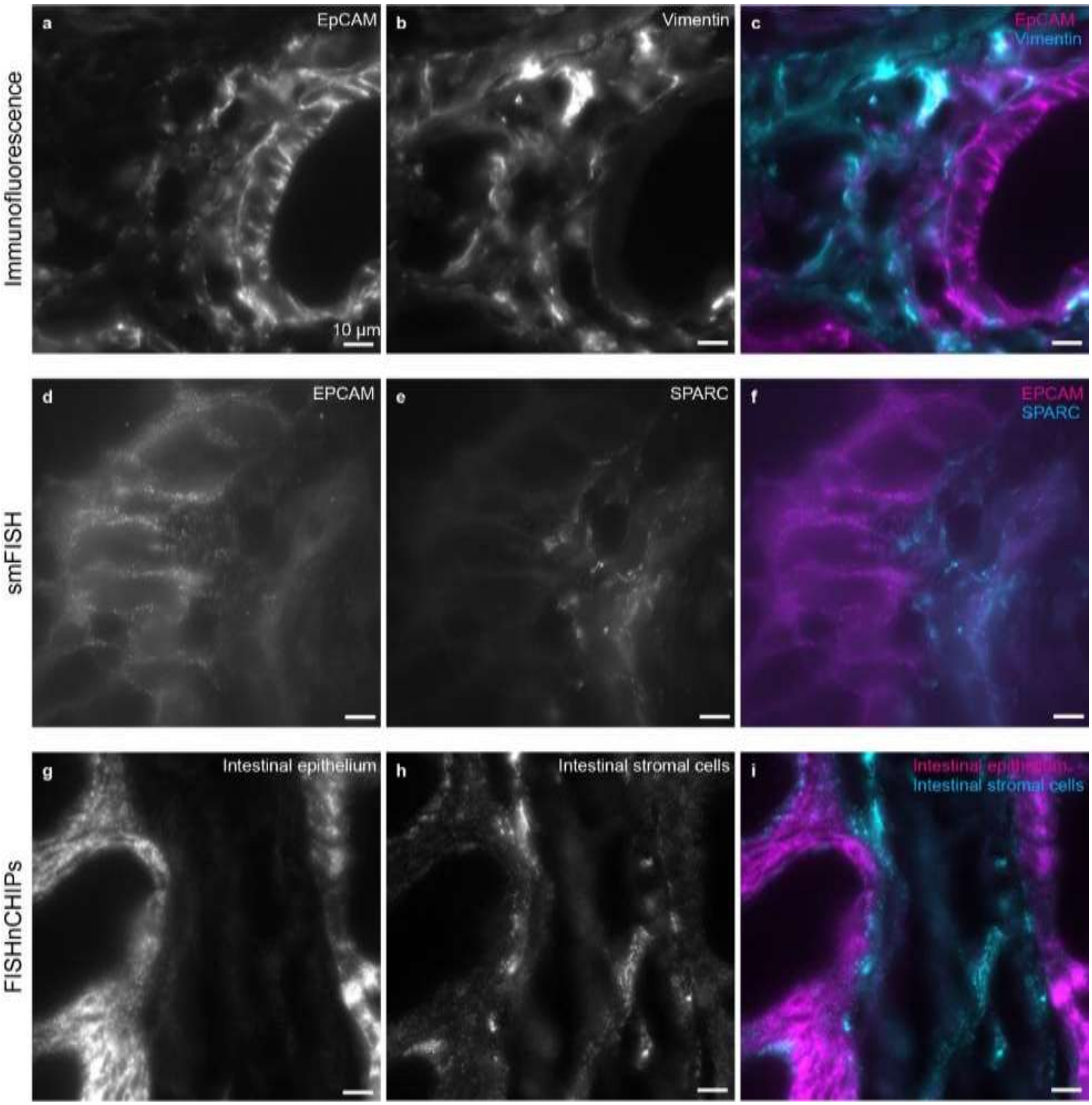

**Extended Data Fig. 15 Comparison of FISHnCHIPs with smFISH and Immunofluorescence staining.**

(**a - b**) Immunofluorescence staining with EpCAM and vimentin antibodies. Scale bars, 10 µm. (**c**) Composite image of immunofluorescence staining: EpCAM (magenta) and vimentin (cyan). Scale bars, 10 µm. (**d - e**) smFISH staining of EPCAM and SPARC transcripts. Scale bars, 10 µm. (**f**) Composite image of smFISH staining: EPCAM (magenta) and SPARC (cyan) Scale bars, 10 µm. (**g - h**) FISHnCHIPs staining of intestinal epithelium and stromal cells. Scale bars, 10 µm. (**i**) Composite image of FISNnCHIPs staining: intestinal epithelium (magenta) and intestinal stromal cells (cyan). Scale bars, 10 µm.

### Extended Data Figure 16

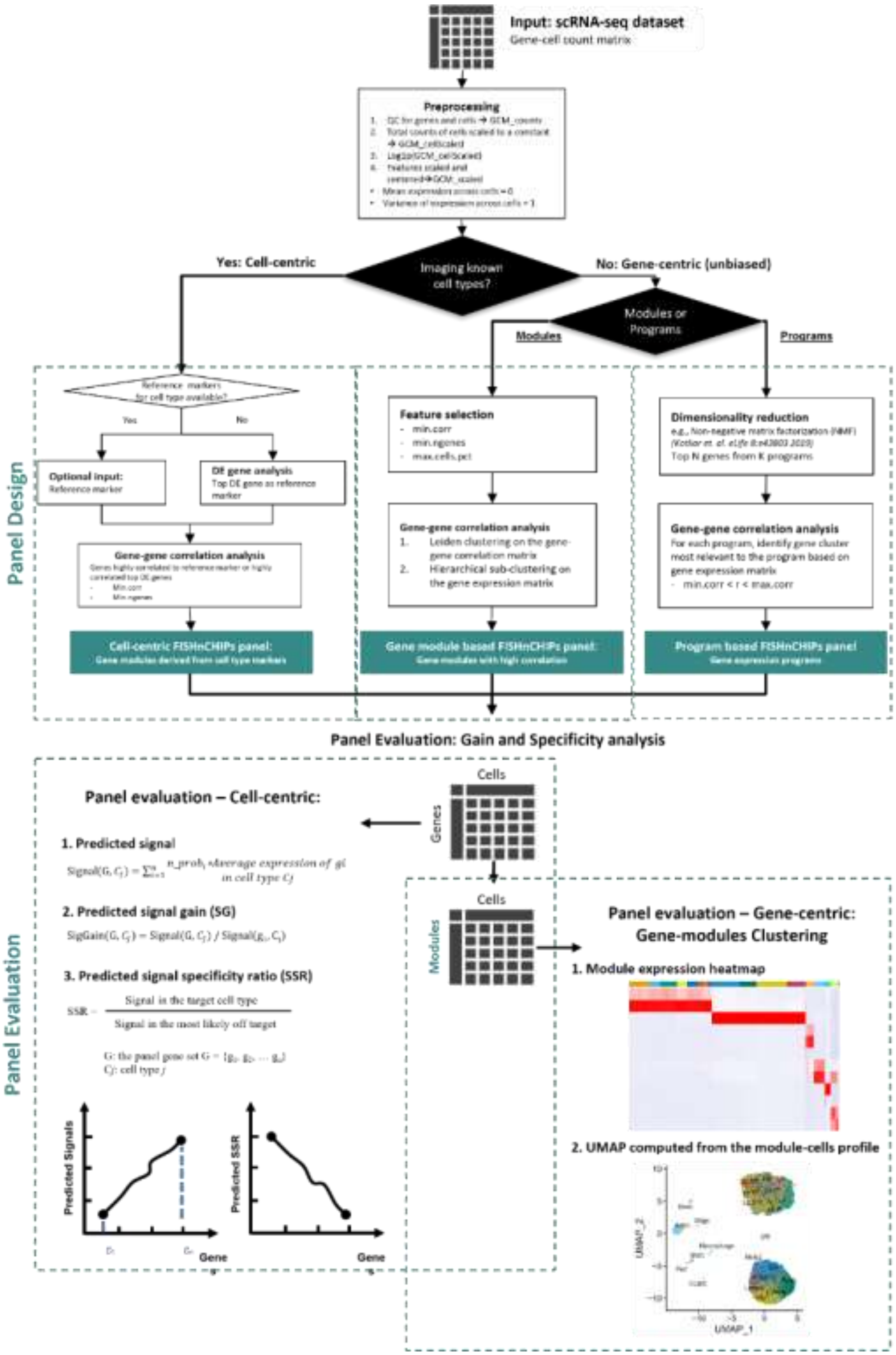


**Extended Data Fig.16 Workflow summarizing the FISHnCHIPs panel design software.**

**Supplementary Table 1. FISHnCHIPs libraries and readout probes.**

The first column is the gene name, the second column is the transcript ID, and the followings columns include readout ID corresponding to the readout probe sequences. All the probe sequences include the forward and reverse primer sequences used during enzymatic library amplification. Related to Figure 2 to 6. Excel spreadsheets contain 6 tabs: “Fig. 2 probe library”, “Fig. 3 probe library”, “Fig. 4 probe library”, “Fig. 5 probe library”, “Fig. 6 probe library”, and “Readout probes”.

Sup Table 1 - FISHnCHIPs_ProbeSequences.xlsx

**Supplementary Table 2. FISHnCHIPs gene panels.**

Gene names and Transcript IDs for all the gene panels. Related to Fig. 2 to 5. Excel spreadsheets contain 5 tabs: “Fig. 2 Gene panel”, “Fig. 3 Gene panel”, “Fig. 4 Gene panel”, “Fig. 5 Gene panel”, and “Fig. 6 Gene panel”.

Sup Table 2 - FISHnCHIPs_Genes.xlsx

**Supplementary Table 3. FISHnCHIPs gene modules and GO terms.**

For the gene modules generated for the mouse brain, gene ontology enrichment analysis was performed to assess the statistical significance of the genes selected. The term source, name, ID, and adjusted *P* values are listed in this table. Related to Fig. 3 and 5. Excel spreadsheets contain 2 tabs: “MouseBrain_Fig3GeneModules” and “MouseBrain_Fig5GeneModules”.

Sup Table 3 - GOTerms_FISHnCHIPsGeneModules.xlsx

**Supplementary Table 4. FISHnCHIPs gene-cell count matrix with annotations.**

FISHnCHIPs expression data with spatial coordinates. Related to Fig. 3, 4, and 5. Excel spreadsheets contain 3 tabs: "FISHnCHIPs_Fig3_data", "FISHnCHIPs_Fig4_data" and "FISHnCHIPs_Fig5_data".

Sup Table 4 - FISHnCHIPsCountMatrix_Annotated.xlsx

**Supplementary Table 5. A comparison of FISHnCHIPs to MERFISH.**

Each of the cell type annotated in MERFISH is tabulated. Related to Fig. 3. Excel spreadsheets contain 1 tab: "ClusterLabelsMap".

Sup Table 5 – MapClusterLabels.xlsx
